# Minor intron containing genes: Achilles’ heel of viruses?

**DOI:** 10.1101/2022.09.30.510319

**Authors:** Stefan Wuchty, Alisa K. White, Anouk M. Olthof, Kyle Drake, Adam J. Hume, Judith Olejnik, Elke Mühlberger, Vanessa Aguiar-Pulido, Rahul N. Kanadia

## Abstract

The pandemic caused by the severe acute respiratory syndrome coronavirus 2 (SARS-CoV-2) revealed the world’s unpreparedness to deal with the emergence of novel pathogenic viruses, pointing to the urgent need to identify targets for broad-spectrum antiviral strategies. Here, we report that proteins encoded by Minor Intron-containing Genes (MIGs) are significantly enriched in datasets of cellular proteins that are leveraged by SARS-CoV-2 and other viruses. Pointing to a general gateway for viruses to tap cellular machinery, MIG-encoded proteins (MIG-Ps) that react to the disruption of the minor spliceosome are most important points of viral attack, suggesting that MIG-Ps may pan-viral drug targets. While contemporary anti-viral drugs shun MIG-Ps, we surprisingly found that anti-cancer drugs that have been repurposed to combat SARS-CoV-2, indeed target MIG-Ps, suggesting that such genes can potentially be tapped to efficiently fight viruses.

## Introduction

As of September 15^th^, 2022, the World Health Organization has reported more than 6.4 million deaths worldwide that were caused by severe acute respiratory syndrome coronavirus 2 (SARS-CoV-2) infection. While future retrospective studies will capture the continuing impact of the current pandemic on long-term human health, the world was undoubtedly not prepared to combat SARS-CoV-2 or any other virus that may cause a future pandemic. Although effective, the development of vaccines against SARS-CoV-2 and other viruses is time-consuming as well as virus-specific, prompting an urgent need to discover and design pan-antiviral therapeutic strategies as a first line of defense (1).

Viruses have co-evolved with their hosts over millions of years, shedding genes that are necessary for their life cycle in favor of a compact genome (2). To recover such lost functions, viral proteins tap the protein-protein interactome of the infected host cells. As a consequence, interactions between host and viral proteins provide insight into the ways a viral pathogen hijacks cellular host function. The first interactome study of SARS-CoV-2 proteins with human host proteins showed that all viral proteins, including the non-structural (Nsp) and structural proteins (nucleocapsid-N, spike-S, membrane-M, envelope-E), interact with the human proteome, providing a promising platform for antiviral strategies (3). Mutations that drive the emergence of new viral strains do not fundamentally alter the replication cycle of the virus, suggesting that virus-host protein interactions can be leveraged to design antiviral drugs. Out of 332 host proteins that appeared in the first interaction map of human and SARS-CoV-2 proteins, 62 were indeed reported drug targets (3). Although these drugs impair the function of individual proteins, viruses often find ways to circumvent such disruptions. This observation points to the need to identify a strategy that simultaneously targets multiple host proteins that are required for various steps of the viral replication cycle. In other words, a unique feature that is shared by multiple host proteins and leveraged by the virus for its replication would be an ideal target to ultimately disrupt virus replication.

Human genes contain both major (>99.5%) and minor (<0.5%) introns, that require the major and the minor spliceosome for their processing. These two types of introns are classified based on the consensus sequences at the 5’ splice sites, branch point sequences, and the 3’ splice sites (4). Because minor introns are often embedded in genes with major introns, splicing of such transcripts relies on the coordinated action of both spliceosomes (5). Splicing of major introns is executed by small nuclear RNAs (snRNA), U1, U2, U4, U5 and U6, and over 150 protein components. In contrast, minor introns are processed by the minor spliceosome that consists of 5 snRNAs (U11, U12, U4atac, U5, U6atac) and many proteins shared with the major spliceosome, as well as 11 unique proteins. Despite the overlap of components between the minor and major spliceosome, splicing of minor introns is mostly carried out by the minor spliceosome to guarantee a fully functional protein (6). Previous studies have shown that disruption of the minor spliceosome, either in human disease or model systems, results in minor intron retention in the MIG transcripts (5, 7), leading to aberrant functions of the resulting MIG-Ps. Specifically, 699 minor intron containing genes (MIGs) exist in the human genome that execute disparate biological functions and are mostly essential for survival, resulting in a high degree of conservation (4, 8).

Here, we explore the role of MIG-encoded proteins (MIG-Ps) that depend on the MiG excision through the minor spliceosome for their expression in viral infections. We show that MIG-Ps are fundamental for the viruses to target host cells and tap cellular machinery through host factors. Furthermore, we provide evidence that MIGs that react to the disruption of the minor spliceosome are most important points of viral attack, allowing us to show that MIG-Ps indeed are bona fide pan-viral drug targets.

## Results

Interrogating the first SARS-CoV-2-host protein-protein interaction (PPI) map by Gordon et al. (3), we surprisingly found that out of the 332 host proteins interacting with SARS-CoV-2 proteins, 20 were MIG-Ps (8) that were involved in every stage of the viral replication cycle (**Fig. 1A**). To test whether these 20 MIG-Ps were in fact significantly enriched, we performed a permutation analysis by randomly sampling 10^5^ sets of 699 known MIG-Ps out of all known human proteins. Notably, we observed that MIG-Ps were, indeed, significantly enriched among the host proteins that interact with SARS-CoV-2 proteins (3, 9, 10) (P = 6.8 ×10^−7^, Fisher’s exact test). Most of these MIG-Ps execute disparate biological roles that SARS-CoV-2 uses to access various biological pathways of the host (8). Given this unique role MIG-Ps play for SARS-CoV-2, we hypothesized that MIG-Ps might also be leveraged by other viruses. Investigating the enrichment of MIG-Ps in viral-host PPIs of other coronaviruses, we observed significant enrichment of MIG-Ps in human-viral interactomes of SARS-CoV-1 and MERS-CoV through our permutation analysis (**Fig. S1**, P < 3×10^−3^, Fisher’s exact test) (11). Broadening our search to viruses for which large-scale protein interaction data are available, we observed significant enrichment of MIG-Ps in the viral-host PPIs of Zika virus (ZIKV), human immunodeficiency virus (HIV-1), human papillomavirus (HPV-1), influenza A virus (IAV), Epstein-Barr virus (EBV), Ebola virus (EBOV), herpes simplex virus 1 (HSV-1), and hepatitis C virus (HCV) (12) (**Fig. 1B**, P < 0.01, Fisher’s exact test). As a comparison, we tested if a random selection of proteins encoded by genes that are alternatively spliced by the major spliceosome (13) would show the same level of enrichment. Notably, we found that MIG-Ps were significantly more enriched in all analyzed viral-host interactomes compared to alternatively spliced genes (**Fig. 1B**, P < 10^−10^, Student’s test).

**Figure 1.**
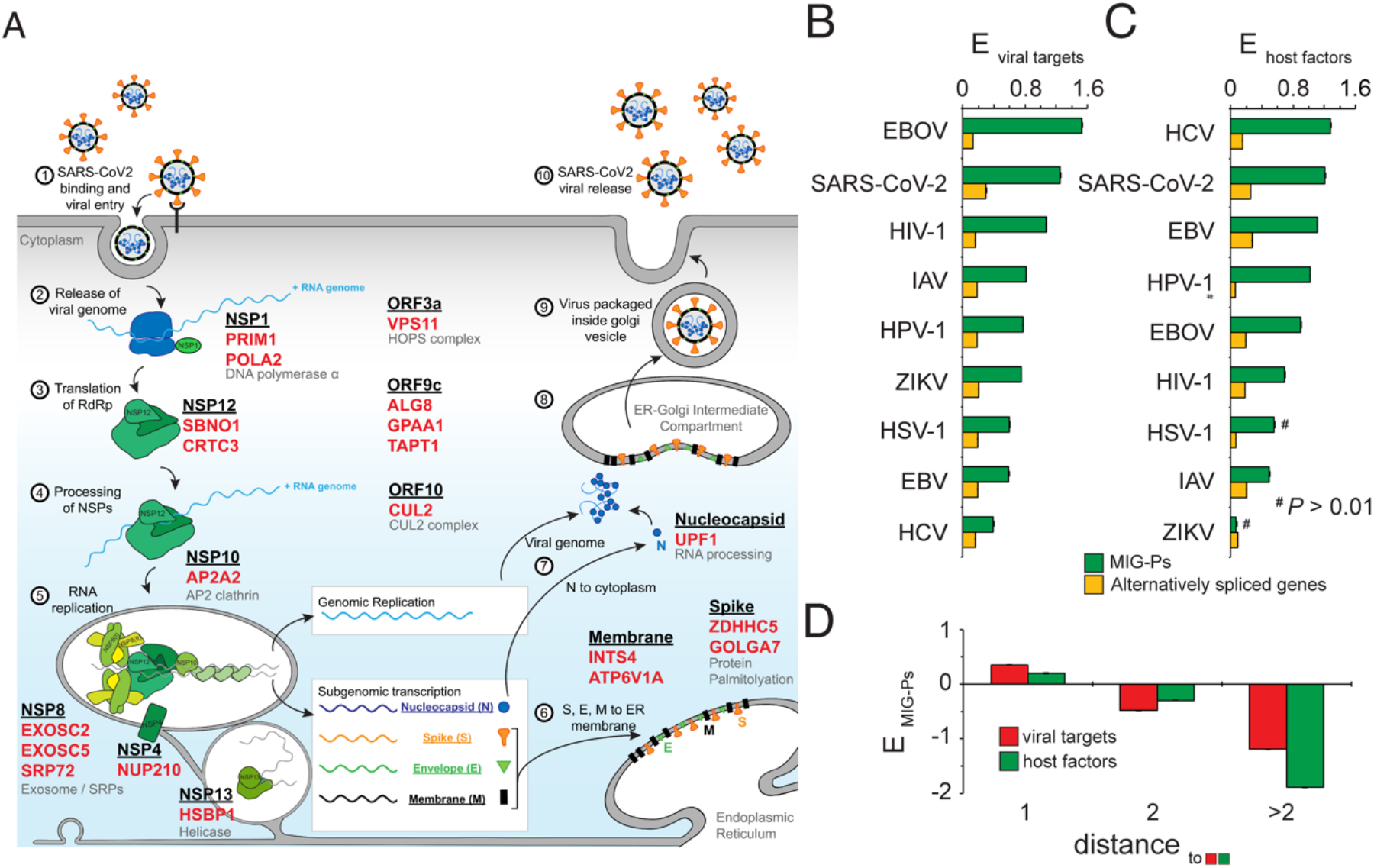
Viral targets and host factors are enriched with MIG-encoded Proteins (MIG-Ps). (*A*) Scheme of the SARS-CoV-2 replication cycle indicating viral proteins (black, underlined) that target MIG-Ps (red) as initially reported in (3). (*B*) Randomly permuting MIG-Ps in comparison to proteins with alternatively spliced isoforms, we observed that MIG-Ps were significantly more enriched (E) in sets of human proteins that were targeted by a variety of viruses. Error bars refer to 95% confidence interval. Enrichment score comparisons were all significant (P < 0.01). (*C*) Similarly, MIG-Ps were enriched in sets of host factor genes that are required by different viruses to infect their host cells (P < 0.01). (*D*) Furthermore, we determined the distance of each protein in a network of human PPIs to the nearest viral target or host factor gene. Randomly sampling MIG-Ps, we observed that MIG-Ps were significantly enriched in the immediate network vicinity of viral targets and host factors and are depleted with increasing distance (P < 0.01).

The significant enrichment of MIG-Ps in virus-host protein interactions led us to interrogate whether MIG-Ps are enriched as viral host factors, defined as host proteins that do not necessarily interact with viral proteins but can either facilitate virus replication (proviral activity) or block virus replication (antiviral activity). Loss of a proviral host factor would block virus replication, whereas loss of an antiviral host factor would enhance virus replication (12). We found significant (P < 0.01) enrichment of MIG-Ps in a combined set of pro- and antiviral host factors for SARS-CoV-2 (14-19) as well as HIV-1, HPV-1, IAV, EBV, EBOV, and HCV (12), while we did not find significant enrichment for ZIKV and HSV-1 (**Fig. 1C**). As previously observed, enrichment of MIG-Ps among host factors was notably stronger compared to genes that were alternatively spliced by the major spliceosome (**Fig. 1C**, P < 10^−10^, Student’s test), suggesting that MIG-Ps execute functions that are exploited by viruses for their propagation cycle.

Although MIG-Ps significantly interact with viral proteins (*i.e*. viral targets) and are host factors, most of the viral targets and host factors are in fact non-MIG-Ps. Therefore, we explored whether MIG-Ps are also part of the molecular network downstream of such viral targets and host factors. In particular, we considered a network of human protein-protein interactions (20) and calculated the shortest distance of each human protein to the nearest viral target or host factor. As a result, proteins that directly interact with a viral target or host factor are at a distance of *d*=1, while a protein that interacts through an intermediary is at a distance of *d*=2. To determine if MIG-Ps are enriched in these groups of proteins that are at a given distance away from a viral target or host factor in the underlying human protein interaction network, we performed a perturbation analysis using SARS-CoV-2 as our initial virus of interest. We randomly sampled 10^5^ sets of MIG-Ps from all human proteins in the underlying interaction network and discovered that MIG-Ps were significantly enriched at a distance of *d=*1 (P < 10^−3^) and were depleted at further distances (P < 10^−3^, **Fig. 1D**). Since we made the same observations for all viruses separately (**Fig. S2**), we conclude that MIG-Ps may be essential components of the molecular network that a wide variety of viruses leverage to enhance their replication, suggesting that MIG-Ps serve as proviral factors.

Since MIG-Ps are significantly present in sets of targets and host factors of individual viruses, we wanted to explore whether each virus leverages distinct sets of MIG-Ps or a core set of MIG-Ps. Therefore, we grouped human proteins into bins based on the number of viruses that use them as target proteins or host factors. Performing a perturbation analysis by randomly sampling sets of 10^5^ MIG-Ps, we, indeed, found that MIG-Ps were increasingly enriched in bins of proteins targeted by multiple viruses (P < 0.01, **Fig. S3A**). As a corollary, we determined the largest number of viruses that leverage pairs of MIG-P viral targets and host factors (**Fig. S3B**). Through a cluster analysis, we found an island that represents those MIG-Ps that are utilized by the greatest number of viruses (right inset, **Fig. S3B**). Such an assembly of MIG-P viral targets and host factors might indicate a molecular framework that different viruses leverage through their co-evolution with the host. In other words, viruses shed genes for functions that are essential and executed through evolutionarily conserved MIG-Ps. To investigate this hypothesis, we collected all genes from 342 cell lines that are essential for cell survival (21). These essential MIGs (*i.e*. essentialome) are largely evolutionary conserved and often date back to the last eukaryotic common ancestor (8). Notably, the vertical sidebar in our heatmap in **Fig. S3B** suggests that MIG-Ps leveraged by many viruses were, indeed, significantly essential for survival (P < 1.7×10^−6^, Fisher’s exact test).

Based on this qualitative observation, we hypothesize that viruses utilize functions of MIG-Ps that remain relatively unchanged across evolution as a backbone that can be stably exploited. Indeed, we observed that MIG-P viral targets of various viruses were enriched with genes in the essentialome (**Fig. S4**), exceeding enrichment signals we obtained by just considering MIG-Ps in general. Furthermore, we obtained similar results when analyzing the enrichment of MIG-P host factors in the essentialome (**Fig. S4**), confirming that conserved MIG-P viral targets and host factors might provide essential host cell functions viruses can reliably bank on to secure viral replication.

The unique position MIG-Ps hold for virus replication suggests that expression of MIG-transcript levels would remain unaltered upon virus infection to facilitate a stable viral infection environment. Examining the transcriptomic profiles in SARS-CoV-2, MERS-CoV, and SARS-CoV-1 infections (22), we found that (essential, targeted) MIG-Ps in the corresponding virus-host interactomes were mostly unchanged (*i.e*. non-differentially expressed) over the course of infection (**Fig. S5**). Specifically, we observed that none of the MIG-Ps in the SARS-CoV-2 host interactome were differentially expressed, further bolstering the idea that MIG-Ps play a crucial role in virus replication.

Viruses typically target host proteins that are involved in a high number of interactions with other host proteins (*i.e*. hubs) (23, 24). Generally, these hubs are essential for cell survival and are evolutionarily conserved (25). As MIG-Ps play a fundamental role for viruses, we hypothesized that they would be significantly enriched in bins of highly interacting proteins in our underlying network of human protein interactions. We confirmed our hypothesis through a permutation analysis (**Fig. 2A)** and further observed that this trend was reinforced when we considered essential MIG-Ps. As a corollary, we hypothesized that viruses exploit such topological characteristics of highly interacting proteins through targeting MIG-Ps. Notably, we confirmed that essential MIG-P viral targets were further enriched in bins of highly interacting human proteins as well as host factors (**Fig. S6**), suggesting that simultaneous targeting of MIG-Ps might be a promising pan-antiviral strategy.

**Figure 2.**
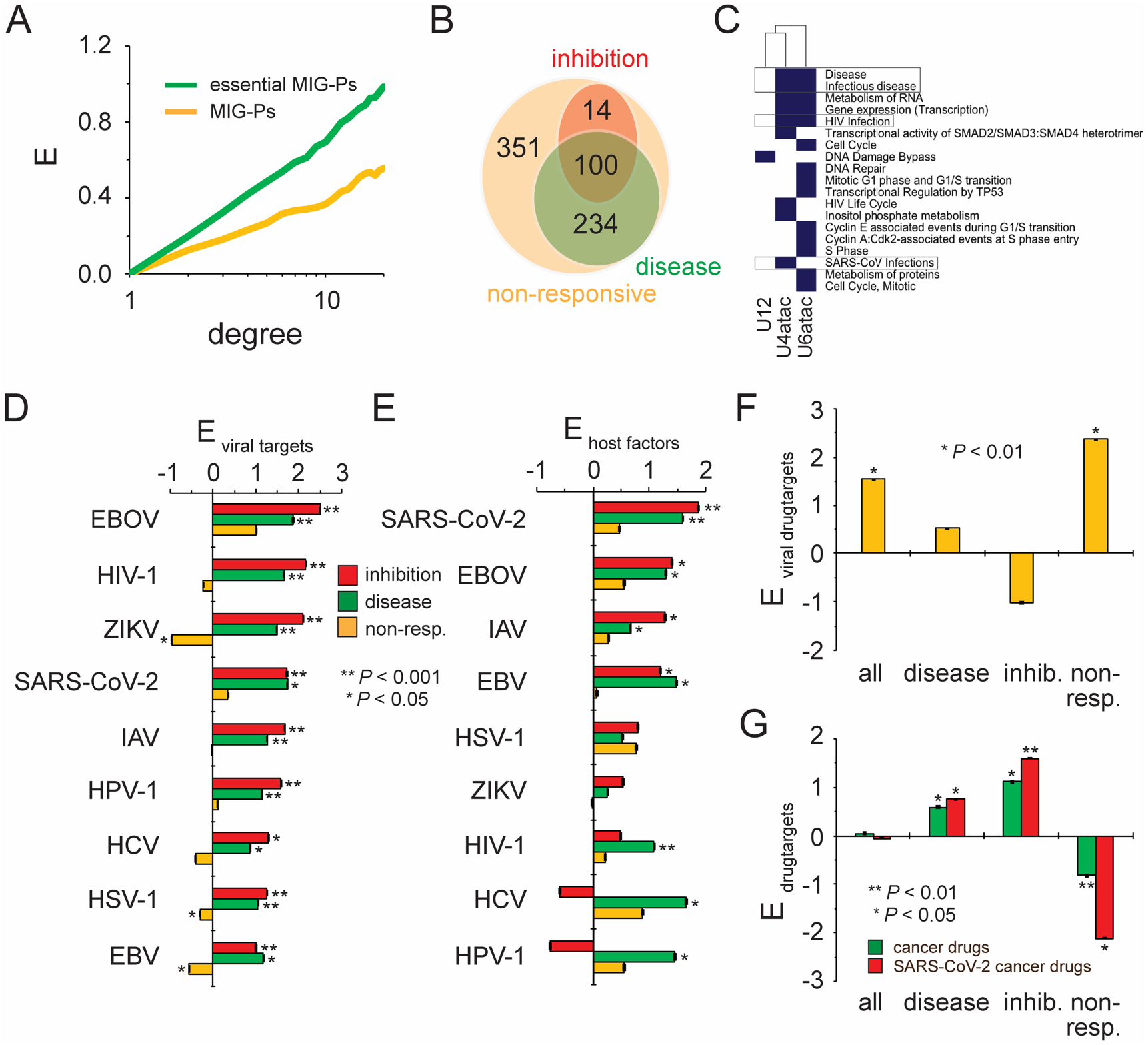
MIG-Ps that are responsive to the disruption of the minor spliceosome are drug targets to combat cancer and viruses. (*A*) In a network of human protein interaction network MIG-Ps are enriched in bins of highly interacting proteins (i.e., higher degree), a trend that is enforced with essential MIG-Ps. (*B*) In the Venn diagram we found a large overlap of sets of MIGs that were responsive to disease mutations and inhibition of the minor spliceosome. (*C*) Considering MIGs that are responsive to minor spliceosome inhibition through morpholinos, targeting the U12, U4atac, and U6atac RNA components, we found a significant enrichment of pathways that correspond to viral infection processes (FDR < 0.05). (*D*) As a corollary, we found that responsive MIG-Ps that were responsive to inhibition of the minor spliceosome were stronger enriched with viral targets, compared to disease responsive and non-responsive MIG-Ps. (*E*) We obtained a similar, yet mixed picture, when we considered enrichment of (non-)responsive MIG-Ps with host factors. (*F*) Considering antiviral drugs, corresponding drug targets were significantly enriched in sets of non-responding MIG-Ps. (*G*) In turn, we found that responsive MIG-Ps were enriched with anti-cancer drug targets. Considering anti-cancer drugs that also combat SARS-CoV-2, we observed an amplification of the trend that drug targets were significantly enriched with MIG-Ps that responded to the disruption of the minor spliceosome.

The observed reliance of viruses on MIG-Ps prompted us to explore whether MIG-Ps could be targeted through minor spliceosome inhibition. To this end, we sought to identify MIG transcripts that respond to minor spliceosome inhibition with either elevated minor intron retention or alternative splicing (7, 26) as a consequence of inhibitor treatment or pathogenic mutations. We obtained 114 responsive MIG-Ps from different datasets including HEK-293T cells treated with morpholinos against U12, U6atac and U4atac snRNA that inhibit the minor spliceosome (**Data S1)**. Among the identified proteins were e.g. DDB1 and PARP1, which also appeared in our cluster of MIG-Ps that were targeted by various viruses in **Fig. S3B**, highlighting the relevance of these responsive MIGs. Focusing on MIGs that were responsive to the disruption of the minor spliceosome through pathogenic variants in spliceosome components (7, 26), we identified 334 MIGs with increased minor intron retention from samples of individuals with Roifman syndrome, Microcephalic Osteodysplastic Primordial Dwarfism Type I and Myelodysplastic syndrome (**Data S1**). Notably, most of MIG-Ps that were responsive to minor intron inhibition were also affected in the context of these diseases (**Fig. 2B**). Next, we analyzed the involvement of MIG-Ps that were responsive to inhibition in cellular functions, allowing us to find pathways that revolved around iviral infection processes (FDR < 0.05, **Fig. 2C**). Focusing on MIG-Ps that responded to disease-based disruption of the minor spliceosome, we also found significant enrichment of mis-spliced MIG transcripts of MIG-Ps in viral infection processes, suggesting that the disruption/inhibition of the minor spliceosome might interfere with viral replication (FDR < 0.05, **Fig. S7**).

The observation that responsive MIGs are enriched in pathways that play a role in viral infections led us to speculate that such genes are exploited by a variety of viruses. Indeed, corresponding MIG-Ps that respond to the inhibition or disease-based disruption of the minor spliceosome are significantly enriched with targets of various viruses, whereas we largely observed the opposite for non-responding MIG-Ps (**Fig. 2D**). Specifically, inhibition-responding MIGs were generally more enriched with viral targets, compared to disease-based responding MIG-Ps (P < 10^−10^, Student’s t-test). We obtained similar results, when we focused on host factors that are required by viruses (**Fig. 2E**). Yet, the enrichment patterns were less uniform compared to viral targets, as host factors of some viruses were more enriched with disease than inhibition responding MIG-Ps.

Since responding MIG-Ps are significantly targeted by viruses to enhance viral infection, we sought to determine whether existing antiviral drugs might target these proteins. We used a comprehensive set of drugs against viruses from the DrugVirus.info databank (27) for analysis. Most of the antiviral drugs in this database have been experimentally tested, and a minority already entered clinical trials or are FDA approved (**Fig. S8**). Performing a permutation analysis through randomly sampling 10^5^ sets of MIG-Ps, we unexpectedly found that antiviral drug targets were enriched with non-responsive MIG-Ps (**Fig. 2F**) instead of responsive MIG-Ps.

Recent reviews of drug repurposing indicated a substantial amount of current clinical trials that assess the efficacy of anti-cancer drugs to combat SARS-CoV-2 (28-30), suggesting that anti-cancer drugs may be targeting MIG-Ps. Therefore, we included 287 licensed cancer drugs from the cancerDrugsDB database (31) in our analysis, 90% of which are FDA approved. Repeating our enrichment analysis, we observed that corresponding drug targets were significantly enriched with responding MIG-Ps, suggesting that these MIG-Ps are *bona fide* targets to combat cancers (**Fig. 2G**). Next, we compiled a list of 29 anti-cancer drugs from (28-30) that are currently tested in clinical trials against SARS-CoV-2 and are included in our set of 287 licensed cancer drugs (**Data S2**). Considering the gene targets of these drugs, we found significant enrichments of MIG-Ps that respond to the disruption of the minor spliceosome through inhibition or mutations, clearly indicating that, indeed, responding MIG-Ps are druggable pan-viral points of intervention.

## Discussion

Our analysis of the involvement of MIG-Ps in viral infections clearly indicates that MIG-Ps in general appear to be commonly exploited by viruses. Our observations suggest that the identified MIG-Ps are significantly targeted through physical interactions between proteins of a variety of viruses including SARS-CoV-2 and the human host cells. Furthermore, MIG-Ps also play a role as host factors that are necessary for virus replication but do not directly interact with viral proteins. Focusing on genes that are essential and strongly evolutionarily conserved, we observed a reinforcement of the original enrichment signal, suggesting that MIGs form a stable backbone that viruses can reliably and efficiently leverage to tap cellular functions for their propagation. Further support for this hypothesis comes from the observation that MIG-Ps are stably expressed upon virus infections, further bolstering the idea that MIG-Ps provide a stable gateway, generally allowing viruses to hijack a host cell.

As the minor spliceosome is needed process MIGs as molecular anchors, allowing viruses to propagate, we hypothesized that MIG-Ps that respond to minor spliceosome inhibition with either elevated minor intron retention or alternative splicing due to inhibitor treatment or pathogenic mutations may point to a pan-viral way forward to combat viruses. Strikingly, we found that inhibition of the minor spliceosome through naturally occurring mutations or application of inhibitors of molecular components of the minor spliceosome mostly hit MIG-Ps that were involved in viral activity and diseases. Furthermore, such responding MIG-Ps were significantly present among viral targets and host factors. Despite their strong involvement in viral-host interactions, contemporary anti-viral drugs largely hit other drug targets than MIG-Ps. do not hit them as drug targets. However, we found responding MIG-Ps enriched with targets of approved anti-cancer drugs. Notably, these signals were enhanced when we focused on targets of anti-cancer drugs that are considered as potential therapeutics against COVID-19, strongly suggesting that targeting MIGs that respond to the inhibition of the minor spliceosome is, indeed, a potential pan-antiviral strategy. In **Fig. S9**, we highlighted drugs that target responding MIG-Ps, suggesting that corresponding drugs affect different pathways. For example, Nivolumab and Pembrolizumab combat the up-regulation of immune checkpoints upon SARS-CoV-2 infection, while Bevacizumab tackles the dysregulated inflammatory response observed in severe COVID-19 (28), and Etoposide blocks DNA synthesis. Furthermore, we found that some of the identified drugs not only target SARS-CoV-2 but also other viruses. For example, Dasatinib is currently tested against SARS-CoV-2, SARS-CoV-1, MERS-CoV, HIV-1, Dengue fever and HCV. Similarly, Gemcitabine shows broad-spectrum antiviral activity against SARS-CoV-2, SARS-CoV-1, HIV-1, HSV-1, and ZIKV (27), strongly suggesting that targeting responsive MIGs is indeed a viable pan-viral therapeutic options. Our observations suggest that although designed with different mechanisms in mind, anti-cancer drugs that also target viruses inadvertently hit responding MIG-Ps. As a corollary, a different way to garner an effect is the therapeutic disruption of the minor spliceosome per se. Notably, treatment with an siRNA that targets the U6atac component of the minor spliceosome led to a lower tumor burden of prostate cancer compared to the current state-of-the-art combination therapy (32). Furthermore, a recent clinical trial involving the steroid dexamethasone showed promise in treating severely ill COVID-19 patients, that reduces the expression of small nuclear ribonucleoprotein U11/U12 subunit 35 (snRNP35), a major component of the minor spliceosome (33), indicating that the inhibition of the minor spliceosome may be a rewarding, yet unexplored broad-spectrum antiviral strategy.

## Materials and Methods

### Determination of the essentialome

Meyers et al. (21) identified genes that are essential for the survival of 342 rapidly dividing cancer cell lines correcting for gene copy numbers per cell line, employing a genome wide CRISPR/Cas9 screen. In accordance with recommendations from the Broad Institute, we removed the PK59_PANCREAS cell line from our analysis, since it had failed subsequent quality controls (34). Classification of the remaining 341 cell lines by cancer origin/type was performed based on the cell line data provided in Supplementary Table 1 by Meyers et al. (21). Based on their thresholding we identified 4,360 genes in the essentialome (8).

### Enrichment analysis

Considering proteins with a certain characteristic *d* (e.g., being a certain distance away from a reference protein or having a certain number of interaction partners in a human protein-protein interaction network), we calculated the fraction of proteins that had a feature *i* in each group *d, f*_*i*_(*d*). As a null model, we randomly sampled protein sets with feature *i* of the same size 10,000 times and calculated the corresponding random fraction, *f*_i,*r*_(*d*). The enrichment/depletion of proteins with feature *i* in a group *d* was then defined as *E*_*i*_(*d*) = lg_2_[*f*_*i*_(*d*)/*f*_i,*r*_(*d*)].

### Pathway enrichment analysis

We utilized the DAVID tool (35) to find enrichments of gene sets in Reactome pathways (36). In particular, we applied hypergeometric tests and corrected P-values using the Benjamini-Hochberg correction (37).

### Human interaction networks

As for a set of high-quality human protein-protein interactions we utilized 165,058 interactions between 15,619 human proteins as of the HINT database (38).

### Human-viral protein interactions

We collected 676 human proteins that interacted with proteins of SARS-CoV-2 from (3, 9, 10). Furthermore, we used 365 targets of SARS-CoV-1 and 292 targets of MERS-CoV from (11). As for other viruses, we collected 823 targets of ZIKV, 1,709 of HIV-1, 2,126 of HPV, 2,797 of IAV, 1,141 of EBV, 332 of EBOV, 2,197 of HSV-1, and 1,033 of HCV as of the HVIDB database (12).

### Viral host factors

We collected 799 human host factors of SARS-CoV-2 from (14-19) as well as 917 in HIV-1, 790 in ZIKV, 315 in HPV, 1,251 in IAV, 144 in EBV, 377 in EBOV, and 262 in HCV as of the HVIDB database (12).

### Alternatively spliced genes

We collected 13,595 genes that have alternative spliced isoforms from the VastDB database (13).

### Antiviral drugs

We tapped the DrugVirus.info databank (27) and downloaded all drugs for the treatment of virus infections in general, including cell cultures, primary cells organoids, animal model, drugs in the different clinical testing phases as well as approved drugs and their corresponding protein/gene drug targets. In particular, we found 42 drugs to treat SARS-CoV-2 that apply to 182 human drug targets, while we obtained 30 drugs and 130 targets for the treatment of HIV-1, 30 drugs and 130 targets for EBOV, 44 drugs and 223 targets for HCV, 31 drugs and 192 targets for HSV-1, 48 drugs and 231 for IAV, 28 drugs and 141 drugs for MERS-CoV and 35 drugs and 167 targets of ZIKV.

### Anti-cancer drugs

We collected 287 licensed cancer drugs used in the treatment of cancer from the cancerDrugsDB database (31), where source data comes from the NCI, FDA, EMA and other data sources. In particular, each drug is approved by official US, European or worldwide agencies. Drugs which are used in cancer treatments to alleviate symptoms or other supportive care uses or which are used for diagnostic purposes are not included. Investigational agents and experimental treatments being used in clinical trials are also not included (31).

### Gene expression

Data from samples infected with three different coronaviruses, including SARS-CoV-2 (22) (GSE150316), MERS-CoV and SARS-CoV-1 (GSE56192), were analyzed for changes in gene expression. Raw counts from RNAseq data pertaining to lung biopsies of SARS-CoV-2 patients and healthy controls were downloaded from SRA. DESeq2 (39) was utilized to process the data, applying variance stabilizing transformation (VST), and to calculate differential expression. Outliers were removed based on principal component analysis (PCA).

Log2 fold changes and adjusted p-values were retrieved for those minor intron containing genes (MIGs) within the SARS-CoV-2 interactome (3) to determine whether any of the MIGs were differentially expressed. Additionally, DESeq2-processed data was downloaded for MERS-CoV and SARS-CoV-1 (40). Data pertaining to MRC5 cell lines 24h post-infection at a multiplicity of infection (MOI) of 3 was included in the current analysis. Log2 fold changes and adjusted p-values were retrieved for MIGs targeted by numerous viruses.

### Minor intron retention and alternative splicing around minor introns

Retention and alternative splicing of minor introns was evaluated in three RNAseq datasets where the minor spliceosome was inhibited (7, 26) (GSE96616). Reads were aligned to the mm10 or hg38 genome using Hisat2, followed by extraction of the uniquely mapped reads (41). The level of minor intron retention and alternative splicing around minor introns was then reported as a mis-splicing index (MSI) as described previously (4). Briefly, for minor intron retention, exon-minor intron boundary reads were quantified using BEDTools and normalized to the number of spliced reads mapping to the canonical exon-exon junction (42). For alternative splicing, spliced reads supporting alternative junctions around the minor intron were quantified and normalized to the number of spliced reads mapping to the canonical exon-exon junction. Significant differences in MSI values between control and minor spliceosome-inhibited samples were determined using a Student’s t-test (U11 cKO) or one-way ANOVA (morpholinos). Since no statistical analyses could be performed for patient data (n=1), elevated retention or alternative splicing was determined as a >2-fold increase in MSI value.

### Open reading frame analysis

The effect of retention and/or alternative splicing on the open reading frame of MIGs was determined using ExPASy (43). To determine whether a premature stop codon would activate nonsense mediated decay (NMD), the distance of the premature stop codon to a downstream exon-junction complex was evaluated. A distance of >50nt is predicted to activate the NMD pathway, whereas in other instances it would produce a protein (44). The effect of the alternative splicing on protein domains was determined by blasting the aberrant protein sequences on the PFAM database (45).

## Acknowledgments

The authors thank M. Wolfinger, I. Hofacker, A. Emili and A. Yousef for fruitful discussions.

## Notes

### Competing Interest Statement

The authors have declared no competing interest.

