## Supplemental Figures for "Minor intron containing genes: Achilles’ heel of viruses?"

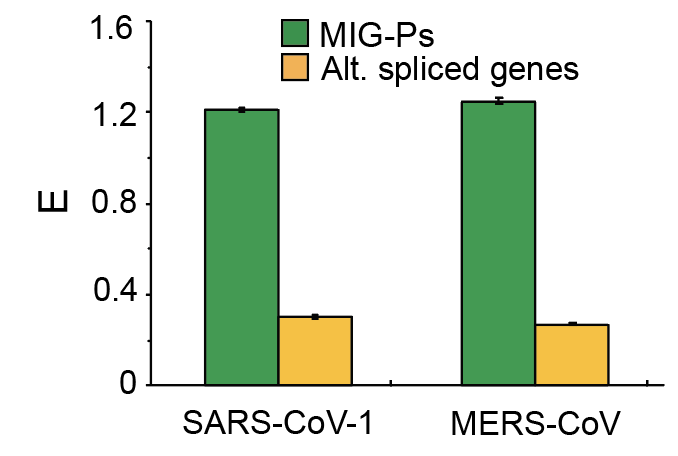

Fig. S1. Viral Targets of SARS-CoV-1 and MERS-CoV are enriched with MIG-encoded Proteins (MIG-Ps). MIG-Ps are enriched (E) in sets of human proteins that interact with SARS CoV-1 or MERS-CoV proteins compared to alternatively spliced isoforms (P < 3$\boldsymbol{\times}$10^-3^, Fisher’s exact test).

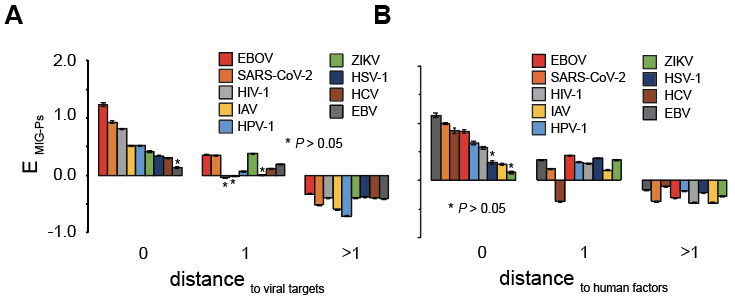

Fig. S2. Distance of each protein in a network of human protein-protein interactions to the nearest viral target *(A)* and host factor gene (B), respectively. Notably, MIG-Ps are enriched (E) in the immediate vicinity of viral targets and host factors.

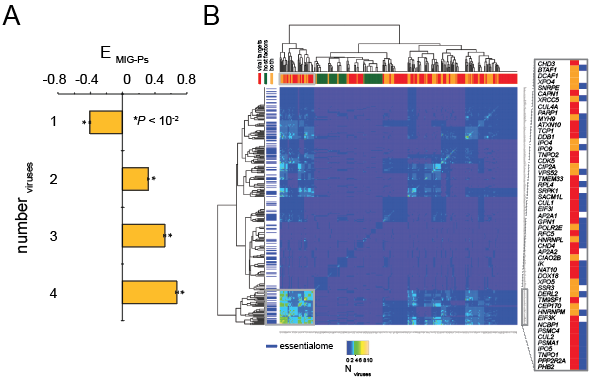

**Fig. S3.** MIG-Ps appear as targets/host factors of many different viruses. In (*A*), we determined the enrichment of MIG-Ps in bins of targets and host factors of an increasing number of viruses, suggesting that some MIG-Ps are targeted by many viruses. (*B*) Determining the largest number

of viruses that leverage pairs of MIG-P viral targets and host factors we found an island of MIG-Ps (grey box on the lower right), where most of these MIGs appeared in the essentialome.

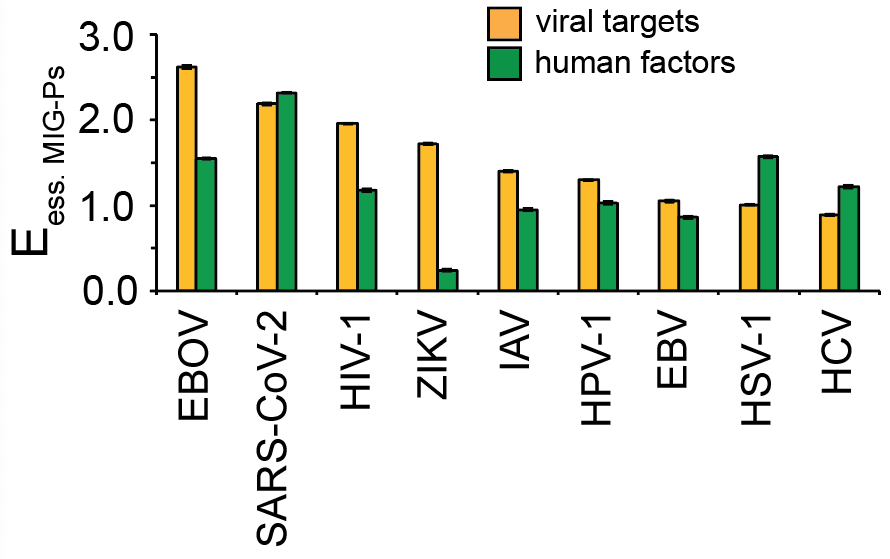

**Fig. S4.** Randomly sampling essential MIG-Ps we found significantly increased enrichment (E) of genes that interacts with viral proteins or were host factors (P < 10^-3^) compared to all MIG-Ps (see Figs. 1B and C in the main paper)

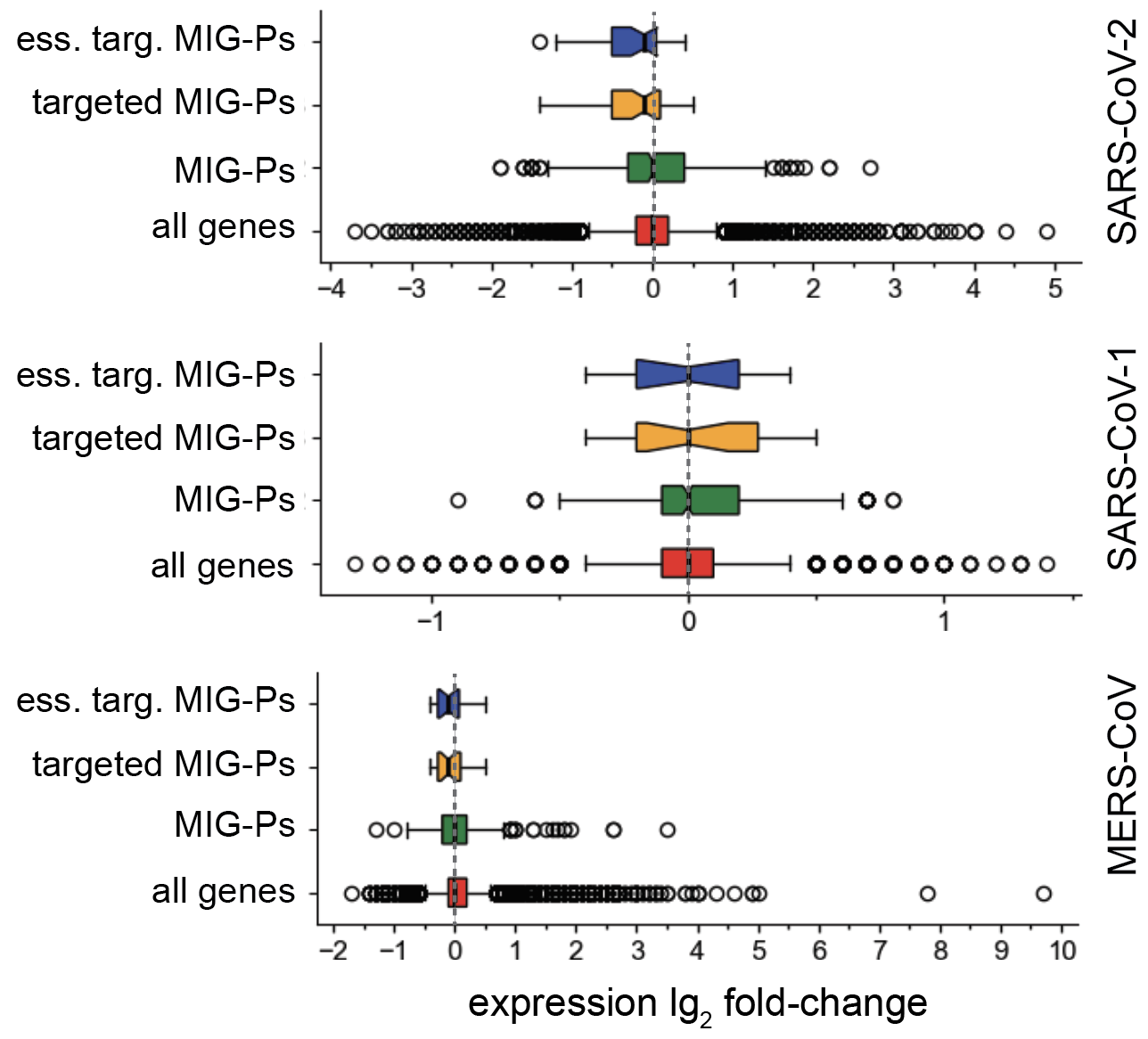

**Fig. S5.** Expression of MIG-Ps upon viral infection. Box-and-whisker plots show that virus-targeted MIG-Ps hardly change their expression values (lg_2_ fold-change) in lung biopsy samples from COVID-19 patients and MRC5 cells infected with MERS-CoV or SARS-CoV-1 at 24 hours post infection.

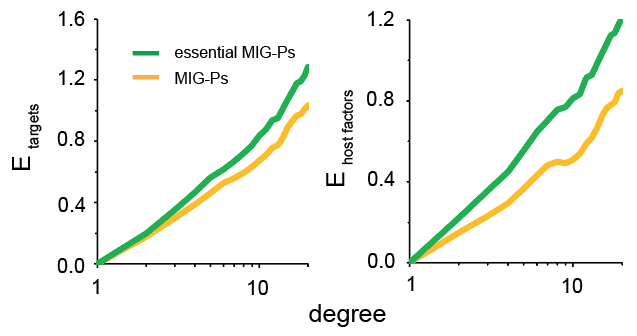
**Fig. S6.** Enrichment of MIG-Ps in a network of human protein interactions. MIG-Ps that are targeted by viruses are predominantly enriched (E) in bins in highly connected proteins in a network of human protein interactions. These trends are reinforced for evolutionarily conserved MIG-Ps. Refining our analysis, we also considered the enrichment of human factors in bins of increasingly connected human proteins.

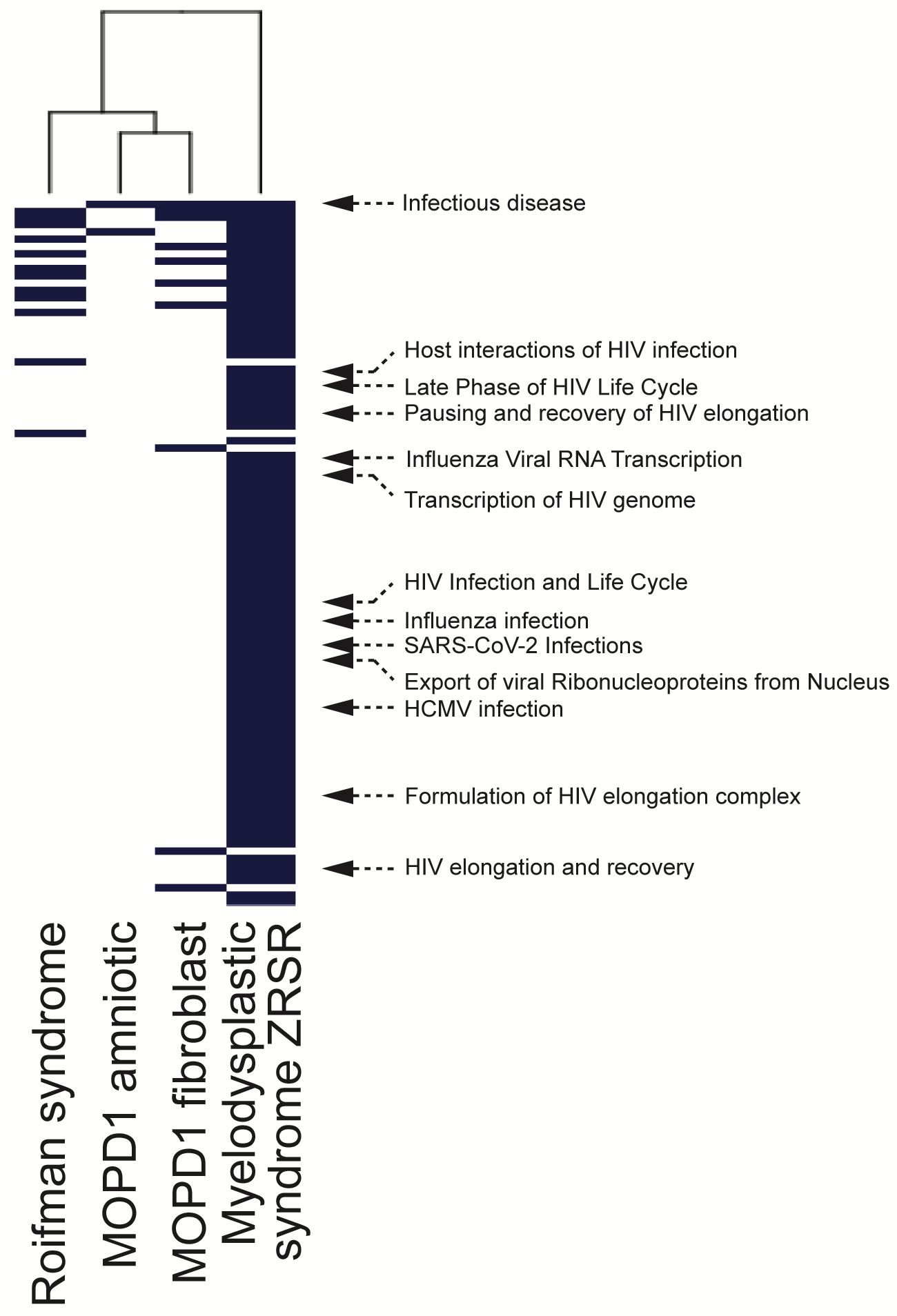

**Fig. S7.** MIGs that were responsive because of mutations in minor spliceosome components in patients that suffer from Roifman, Microcephalic Osteodysplastic Primordial Dwarfism, Type I (MOPD1) or Myelodysplastic syndromes. Reactome Pathway enrichment indicated involvement of responding MIGs in viral processes on a significant basis (FDR < 0.05).

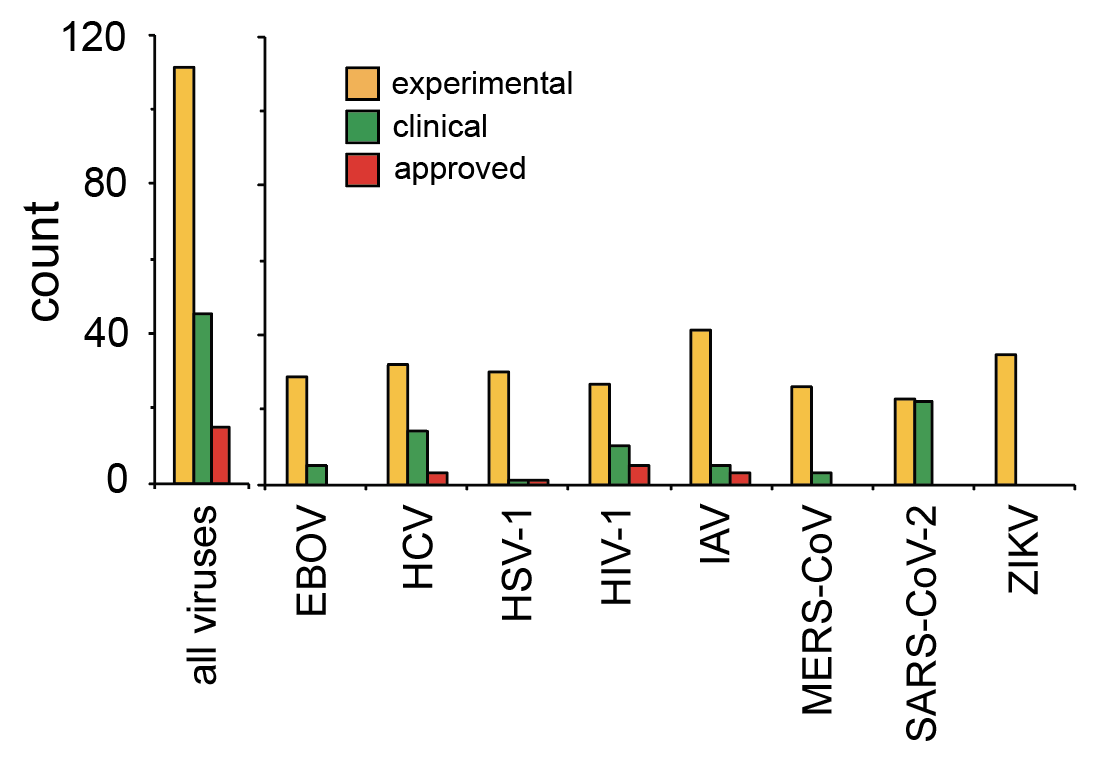

Fig. S8. Stages of testing of all antiviral drugs in the DrugVirus.info database. We found that most of the antiviral drugs have been tested in cell cultures, primary cell organoids and animal models. Some of the drugs are currently tested in clinical trials or are already FDA-approved.

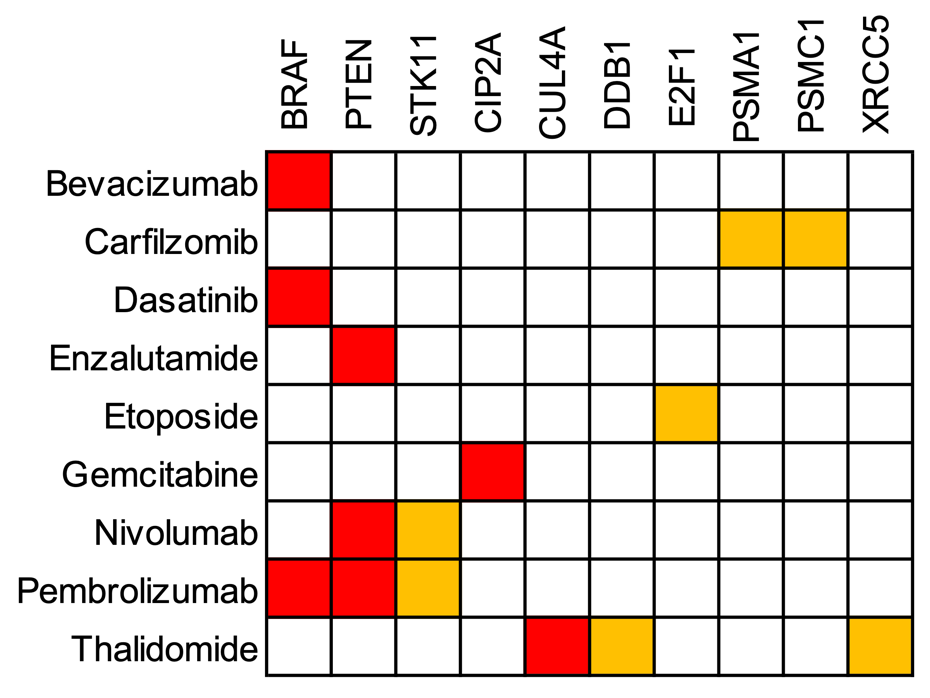

Fig. S9. Matrix of drugs and their drug targets that are responsive MIGs. Red boxes correspond to disease responding MIGs, while orange boxes point to disease and inhibition responding MIGs.

**Dataset S1** (separate file). List of Minor Intron coding Genes that are responsive to minor spliceosome inhibition through morpholinos, targeting the U12, U4atac, and U6atac RNA components and/or naturally occurring mutations/variations in minor spliceosome components in patients that suffer from Roifman, Microcephalic Osteodysplastic Primordial Dwarfism, Type I (MOPD1) or Myelodysplastic syndromes. All minor introns are annotated with their genes where they occur, genomic coordinates, length of minor intron, 5’, 3’ and branch point (BPS) sequences, confidence scores, as well as appearance in diseases and/or inhibition of minor splicesome components as well as the genes responsiveness and occurrence in the essentialome, viral targets, host factors and drug targets.

**Dataset S2** (separate file). List of 29 approved anti-cancer drugs that are/were tested against SARS-CoV-2. Each drug is annotated with agencies that approved the drug (EMA, FDA, EN,WHO), Drugbank ID, ATC identifier, ChEMBL annotation, indication which cancer types are targeted, gene drug targets and clinical trial numbers. Shaded drugs interact with responsive MIGs as drug targets (bold).
